# Structural studies of the Tudor domain from the *Bombyx* homolog of *Drosophila* PAPI: Implication to piRNA biogenesis

**DOI:** 10.1101/786731

**Authors:** Paul A. Hubbard, Xinlei Pan, Randall McNally, Yohei Kirino, Ramachandran Murali

## Abstract

PIWI proteins and their associated PIWI-interacting RNAs (piRNAs) play crucial roles in proper gametogenesis in animal gonads. Partner of PIWIs (PAPI) is one of the important piRNA biogenesis factors. PAPI contains a Tudor domain and tandem KH domains. The tudor domain specifically recognizes symmetrical-dimethylarginines (sDMAs) on PIWI proteins. BmPAPI, a *Bombyx mori* homolog of PAPI, is localized at the outer membrane of mitochondria and supports exonucleolytic trimming of piRNA precursors to form mature 3’-end of piRNAs. To understand the structural basis of piRNA processing by BmPAPI, we present crystal structures of the *apo-* and sDMA-liganded Tudor domain of BmPAPI.

## INTRODUCTION

PIWI proteins are a subfamily of Argonaute (Ago) proteins, which form gene-silencing complexes with PIWI-interacting RNAs (piRNAs). They play important roles in germline development and are well-conserved through evolution (1–4). The PIWI-piRNA machinery protects genomic integrity in animal gonads by silencing transposons, and regulates gene expression to ensure proper gametogenesis (5–7). Disruption of piRNA biogenesis causes germline defects and eventually leads to sterility (8, 9).

PIWI proteins contain four main domains; an N domain, a PAZ domain that recognizes the 3’ end of the interacting RNA, a MID domain that coordinates the 5’ phosphate of the RNA, and a PIWI domain that houses key catalytic residues (10, 11). In addition, symmetrical-dimethylarginines (sDMAs) are conserved in PIWI proteins from various organisms and are present in motifs that contain multiple arginine-glycine or arginine-alanine repeats, typically at the N-termini of PIWI proteins (12, 13). It is reported that arginine methylation is crucial for PIWI-mediated piRNA biogenesis (14, 15).

The Tudor royal family of proteins recognize methylated lysine or arginine residues to mediate protein-protein interactions, and are characterized by the well-studied canonical Tudor core domain structure, which typically comprises 4-5 β-strands that form a β-barrel with an aromatic pocket on the surface to accommodate methylated substrate (16, 17). A subset of the Tudor domain-containing proteins, primarily expressed in the germ cells, binds PIWI proteins through specific interactions with sDMA and is essential in piRNA biogenesis (18–21). PIWI-binding Tudor proteins possess an ‘extended Tudor domain’ with an extra conserved structural element flanking the Tudor core (22, 23). Additionally, in these proteins, the Tudor domain is frequently associated with RNA-binding domains such as the hnRNP K Homology (KH) domain (24–26).

PAPI (Partner of PIWIs) is a Tudor domain-containing protein first discovered in *Drosophila* that specifically interacts with PIWI proteins through recognition of sDMA (27). It plays an essential role in piRNA biogenesis by recruiting PIWI proteins to nuage, the perinuclear granules in *Drosophila* germline cells. Mouse and *Bombyx mori* studies have revealed that disruption of PAPI causes 3’-end extension of piRNAs, suggesting its role in 3’-end maturation of piRNAs (28, 29). Indeed, a recent report showed that BmPAPI, a *Bombyx mori* homolog of *Drosophila* PAPI, supports PNLDC1-catalyzed processing of PIWI-loaded piRNA precursors to form mature 3’-ends of piRNAs (28, 30). BmPAPI contains tandem KH domains and an extended Tudor domain consisting of a core Tudor domain flanking an oligonucleotide-binding fold (OB-fold) domain. The Tudor domain specifically recognizes sDMAs on *Bombyx* PIWI proteins, Siwi and BmAgo3, and drives PIWI-BmPAPI interaction at the outer membrane of mitochondria (28). While the KH domains are expected to interact with piRNAs and/or piRNA precursors, supporting evidence for this is needed and its function in the piRNA biogenesis pathway remains unclear. To better characterize BmPAPI function and the PIWI-PAPI interaction, we initiated the structural studies of BmPAPI. Here we report the crystal structures of the BmPAPI Tudor domain in its unliganded form as well as bound to sDMA. The overall topology of the Tudor domain and the binding site of sDMA is conserved in BmPAPI. An extended loop forms part of the sDMA-binding site which may have functional role in piRNA biogenesis.

## METHODS

### Expression and purification of BmPAPI Tudor domain

The BmPAPI extended Tudor domain region corresponding to Gly245-Gly463 was sub-cloned into pET28a vector to include an N-terminal His-tag and expressed in BL21(DE3). Purification was by IMAC and SEC using a HiLoad 16/60 Superdex column (GE Lifesciences) to produce >95% pure protein as determined by SDS-PAGE analysis.

### Pull-down binding assay

To confirm the ability of the purified BmPAPI Tudor domain to recognize sDMAs of PIWI proteins, stable BmN4 cells expressing Flag-tagged Siwi proteins were utilized for pull-down binding assay as described previously (28). Briefly, the cells expressing wild-type Siwi or sDMA-lacking mutant Siwi, whose sDMAs are replaced with lysines, were cultured at 27°C in Insect-Xpress medium (Lonza). Cell lysates were prepared in a binding buffer containing 50 mM Na-Phosphate (pH 8.0), 300 mM NaCl, 0.01% Tween 20, 1% Triton X-100, and complete protease inhibitor cocktail (Roche Diagnostics). BmPAPI Tudor protein immobilized on Dynabeads His-Tag Isolation and Pulldown (Life Technologies) was incubated with the lysate for 20 min at 4°C and was washed extensively with binding buffer. Bound proteins were then eluted into an elution buffer containing 50 mM sodium phosphate, pH 8.0, 300 mM imidazole, 140 mM NaCl and 0.02% Tween 20. Western blots of the proteins were subsequently performed by using anti-Flag (Sigma Aldrich) for detection of Siwi and anti-His (Cell Signaling) for detection of BmPAPI Tudor.

### Crystal structures of BmPAPI Tudor domain

Protein was concentrated to 10 mg/mL in 20 mM HEPES, pH 7.5 and 150 mM NaCl before being crystallized using the hanging drop method at 18°C. For the *apo*-BmPAPI crystals, the well solution contained 1% PEG 8,000, 50 mM MES, pH 5.5, 100 mM KCl and 10 mM MgCl_2_, and was mixed with protein at an equal volume ratio, with crystals of space group P3221 taking about one week to reach their maximum dimensions of 90 x 90 x 30 μm. Prior to data collection, crystals were dehydrated against air in a sealed chamber for two days before being mounted on nylon loops and cooled to 100 K. To solve the structure of the BmPAPI-sDMA complex, attempts to obtain *apo*-crystals from the previous conditions failed. Thus, a new form of *apo*-BmPAPI crystals (space group P4_3_2_1_2) was obtained using the hanging drop method with a precipitant solution of 0.95 M ammonium sulfate, 50 mM sodium acetate, pH 5.5 and 5% glucose. These crystals were untreated or soaked in 50 mM sDMA salt (Sigma Aldrich) for 3 days before data collection. X-ray diffraction data were collected in-house using a Rigaku MicroMax 007-HF rotating anode equipped with a Rigaku R-AXIS IV++ image plate. BmPAPI structures were solved using the molecular replacement method. The P3_2_21 *apo*-form was determined using human Tudor domain (PDB ID: 3FDR) as the search model. Subsequently, the *apo-* and sDMA-forms of the P4_3_2_1_2 crystals were determined using the *apo*-BmPAPI Tudor domain. Approximately 90% of residues were traced (excluding His-tag), with the models possessing appropriate R-work factors of 20.1%, 25.3%, and 19.5% and R-free factors of 27.7%, 29.6%, and 26.5% for the P3_2_21 *apo*-(PDB ID 5VQG), P4_3_2_1_2 *apo*-(PDB ID 5VY1), and P4_3_2_1_2 sdMA-bound (PDB ID 5VQH) forms, respectively.

## RESULTS AND DISCUSSION

### Tudor domain of BmPAPI specifically binds to sDMAs of PIWI protein

To characterize the Tudor domain of BmPAPI and to confirm its ability to specifically recognize sDMAs of PIWI proteins, we produced and purified His-tagged Tudor recombinant protein and performed pull-down experiments. We immobilized the recombinant Tudor on beads and subsequently incubated the beads with lysates from BmN4 stable cell lines expressing wild-type or mutant Siwi protein. The mutant Siwi protein lacks sDMAs due to substitutions of arginines in the sDMA motif with lysines (Fig. 1A). After extensive washing, the eluates were subjected to western blots to detect the bound Siwi protein. As shown in Fig. 1B, only wild-type Siwi, and not mutant Siwi, was detected in the Tudor-bound fraction, suggesting specific association of the BmPAPI Tudor domain with the sDMAs of Siwi protein.

**Figure 1.**
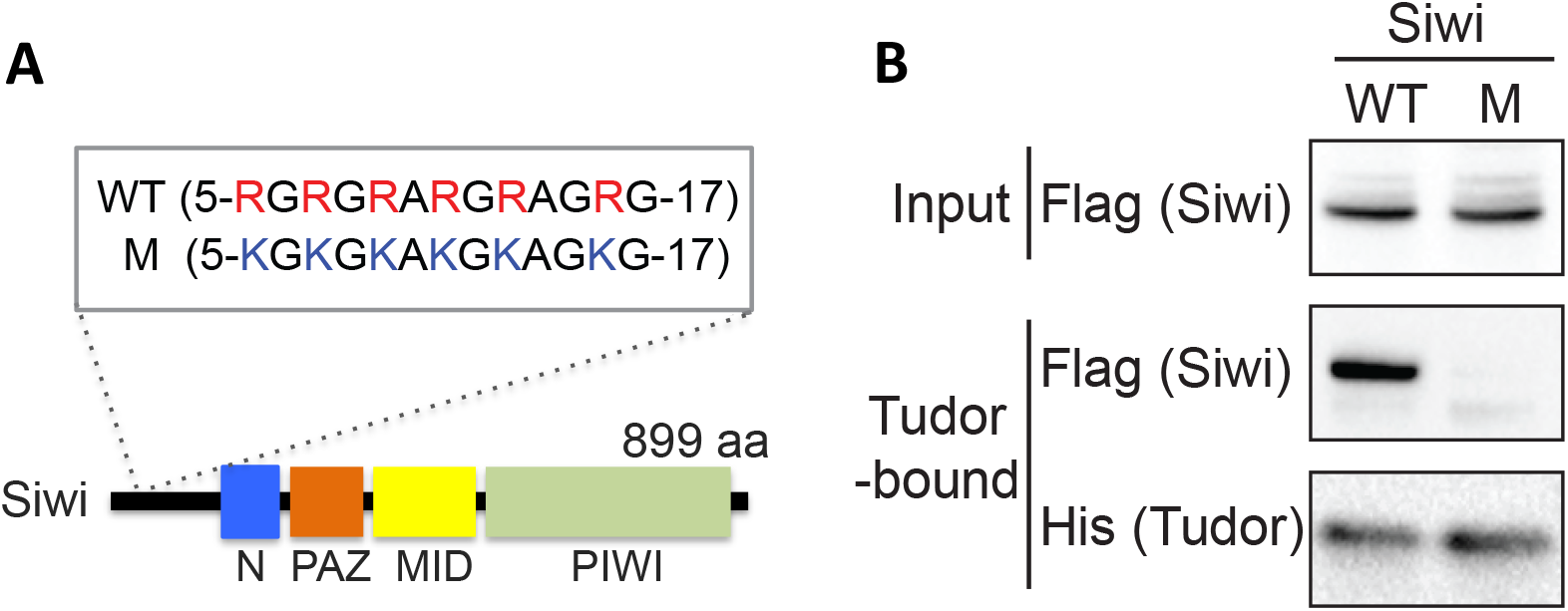
Pull-down experiments to analyze the association of BmPAPI Tudor domain with Siwi protein. **(A)** Domain structure of Siwi wild-type (WT) and mutant (M) Siwi protein sequences showing the arginines in the sDMA motif and their substitution by lysines in the mutant. **(B)** His-Tudor immobilized on Dynabeads was incubated with the lysates from BmN4 cells expressing WT or M Siwi protein. After extensive washing, the eluates were probed with anti-Flag (for Siwi detection) and anti-His antibodies (for Tudor detection).

### Crystal structure of the extended Tudor domain from BmPAPI

To determine the structure of the extended Tudor domain of BmPAPI, crystallization conditions were screened and optimized to yield crystals of space group P3_2_21 that diffracted to 2.6Å (Table 1). The refined *apo*-model contains one protein molecule and 56 water molecules in the asymmetric unit, showing electron density for residues 245-461, excluding flexible loop region residues 379-391, 424-429 and 444-445. The extended Tudor domain of BmPAPI consists of an OB-fold domain and a core Tudor domain (31). The secondary structure comprises 6 α-helices and 10 β-sheets, as is shown in Figure 2. A closed β-barrel is present in the OB-fold domain, comprising anti-parallel β-sheets β1-β2 and β7-β9, and capped by α-helix α5 located between β7 and β8. It has been reported that the OB-fold domain of a *Drosophila* homolog protein was responsible for sensing the position of the sDMA (22). Whether the OB-fold domain of BmPAPI has a similar function is not yet known. The core Tudor domain is formed by four anti-parallel β-sheets, β3-β6. Two long linkers connect the core Tudor domain to the OB-fold domain: helices α1 −α2 at the N-terminus of the core Tudor domain and α3-α4 at the C-terminus.

**Figure 2.**
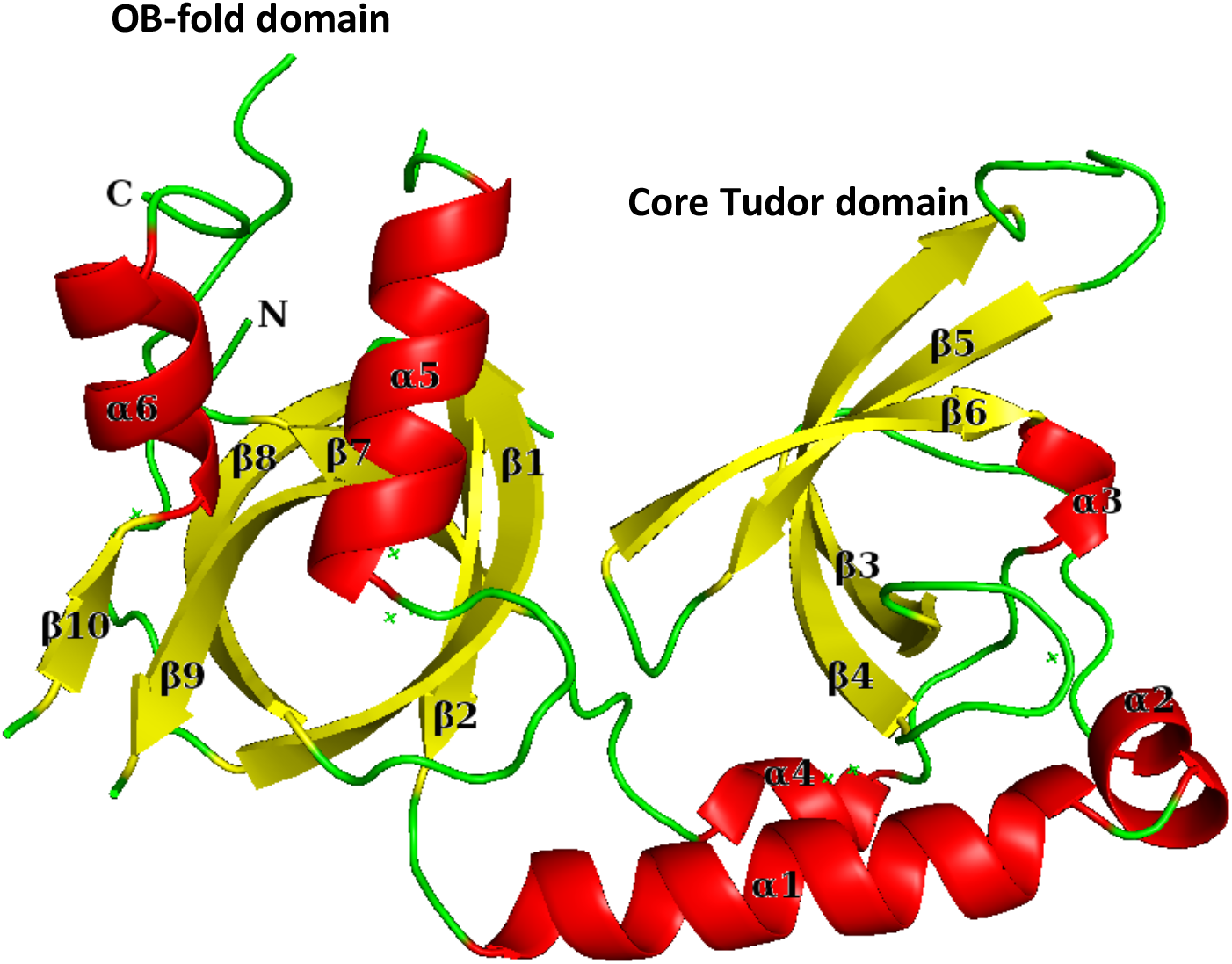
Crystal structure of BmPAPI extended Tudor domain from crystal form P3_2_21. α-helices and β-sheets are labeled and shown in red and yellow respectively. Loops are shown in green. The OB-fold domain is comprised of β1 −β2, β7-β9 and α5. The core Tudor domain consists of β3-β6.

**Table 1.** Data collection and refinement statistics.

| **Data collection** |  |  |  |
| --- | --- | --- | --- |
|  | *apo*-BmPAPI Tudor | sDMA-BmPAPI Tudor | *apo*-BmPAPI Tudor |
| Resolution range (Å) | 31.78 – 2.60 (2.72 – 2.60)<sup>a</sup> | 39.61 – 2.40 (2.51 – 2.40) | 19.50 – 3.05 (3.20 – 3.05) |
| Space group | P3<sub>2</sub>21 | P4<sub>3</sub>2<sub>1</sub>2 | P4<sub>3</sub>2<sub>1</sub>2 |
| Unit cell dimensions *a* x *b* x *c* (Å) | 63.6 x 63.6 x 100.7 | 66.0 x 66.0 x 224.5 | 65.7 x 65.7 x 225.4 |
| Unit cell angles $\alpha$ x $\beta$ x $\gamma$ (°) | 90.0 x 90.0 x 120.0 | 90.0 x 90.0 x 90.0 | 90.0 x 90.0 x 90.0 |
| R<sub>merge</sub> (%) | 10.6 (60.8) | 7.7 (53.5) | 17.1 (57.2) |
| Total number of observed reflections | 31231 (3665) | 68909 (6867) | 26088 (8655) |
| Total number of unique reflections / multiplicity | 7573 (915) / 4.1 (4.0) | 18600 (2275) / 3.7 (3.0) | 8655 (1274) / 3.0 (3.0) |
| Mean I/$\sigma$ | 7.8 (1.6) | 8.7 (1.6) | 5.9 (2.1) |
| Completeness (%) | 99.5 (100.0) | 92.8 (95.0) | 87.9 (90.3) |
| **Refinement** |  |  |  |
| R<sub>work</sub> / R<sub>free</sub> (%) | 20.1 / 27.7 | 19.5 / 26.5 | 25.3 / 29.6 |
| *r.m.s.d.* bond lengths (Å) / angles (°) | 0.011 / 1.416 | 0.012 / 1.482 | 0.002 / 0.654 |
| Number of residues modeled | 195 (1 chain) | 398 (2 chains) | 389 (2 chains) |
| Number of water molecules modeled | 56 | 153 | 0 |
| Average B-factor of all residues (Å<sup>2</sup>) | 34.6 | 45.2 | 51.0 |
| Average B-factor of all waters ($\text{\AA}^2$) | 25.2 | 40.1 | N/A |
| Ramachandran plot (RAMPAGE): |  |  |  |
| Favored | 190 (97.4%) | 378 (95.0%) | 367 (97.9%) |
| Allowed | 5 (2.6%) | 19 (4.8%) | 8 (2.1%) |
| Outliers | 0 (0.0%) | 1 (0.3%) | 0 (0%) |

<sup>a</sup>Values in parentheses are for highest-resolution shell

### Crystal structure of the BmPAPI-sDMA complex

To better understand the interaction between BmPAPI and PIWI proteins, we attempted to solve the structure of sDMA-bound BmPAPI Tudor domain by soaking the *apo-* crystals with sDMA ligand, but were unsuccessful. However, we found crystals of a different space group (P4_3_2_1_2) that were amenable to soaking with sDMA ligand, and solved the structure of the P4_3_2_1_2 form in both *apo*- and sDMA-bound states (Table 1). The structures are solved to 3.05 Å and 2.4 Å (for the *apo*- and sDMA-bound forms, respectively) by molecular replacement using the *apo*-BmPAPI Tudor domain as a search model. The refined sDMA-bound structure contains two protein chains, A and B, and 153 water molecules in the asymmetric unit. Chain A and Chain B have a RMSD of 0.456 Å for 177 Cα atoms aligned (Figure 3A).

**Figure 3.**
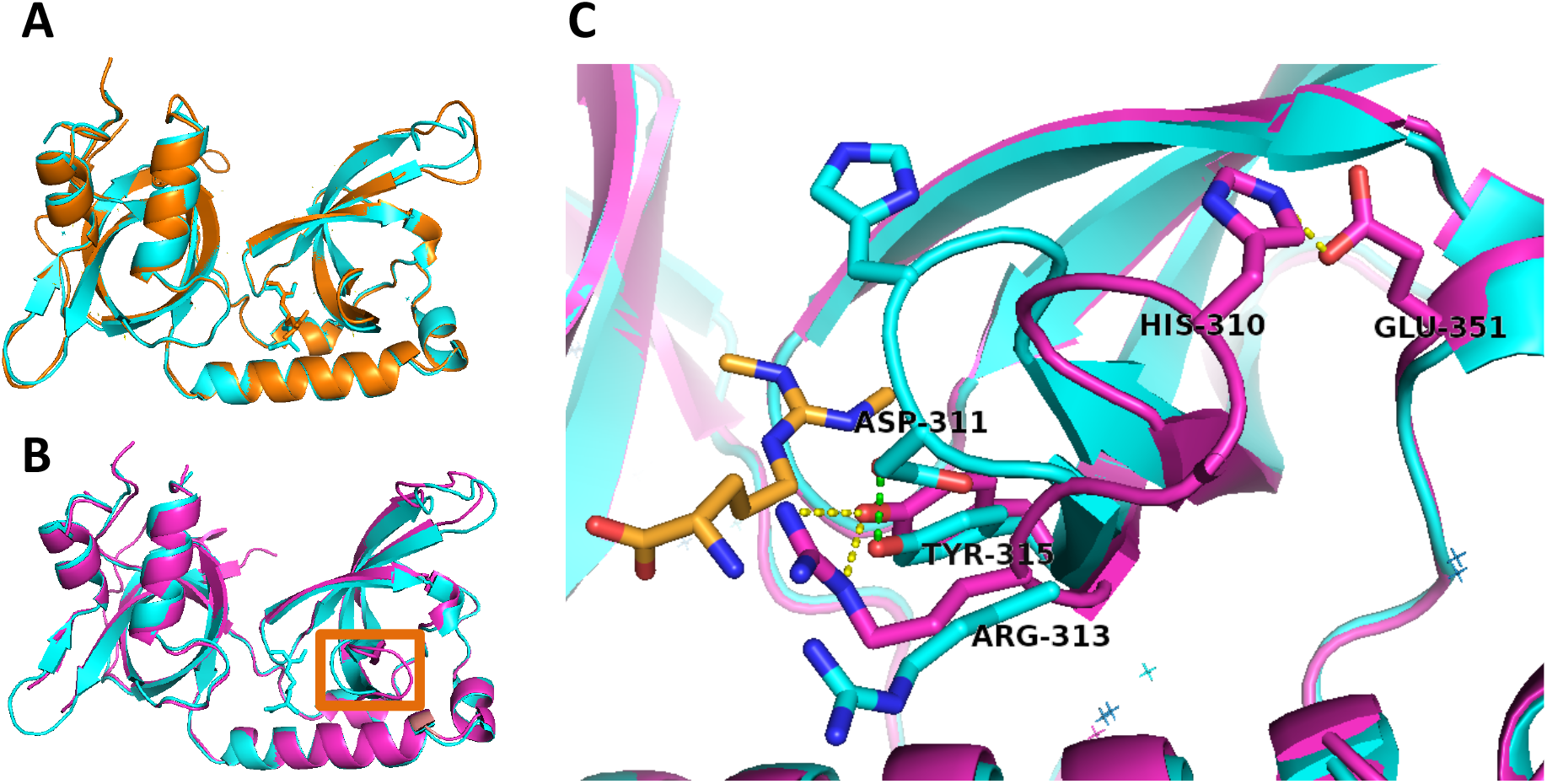
Crystal structure of the BmPAPI-sDMA complex. **(A)** Overlay of the two chains in the BmPAPI-sDMA structure. Chain A is shown in orange and chain B is shown in cyan. **(B)** Superposition of P3_2_21 *apo*-BmPAPI (magenta) and P4_3_2_1_2 sDMA-BmPAPI complex (chain B, cyan) structures. A flexible loop that differs in conformation between the crystal forms is highlighted by the orange box. **(C)** A close-up view of the flexible loop region in native (magenta) and sDMA-bound (cyan) state. The sDMA is shown in orange. Blue and red indicate nitrogen and oxygen atoms respectively. Hydrogen bonds are shown as dashed lines (yellow for the native state and green for sDMA-bound state).

The overall structures of the P3_2_21 *apo*-BmPAPI and P4_3_2_1_2 sDMA-BmPAPI complex are highly similar; the RMSD for backbone structures of *apo*-BmPAPI and sDMA-BmPAPI (Chain B) is 0. 610 Å for 177 Cα atoms aligned. However, comparison of the two structures revealed that the loop consisting of residues 308-314 and inserted between β-strands β3 and β4 has moved in the sDMA-bound form to enclose the sDMA inside the aromatic cage (Figure 3B). A closer view of the loop region shows that in the *apo*-state (open position), Tyr315 is double hydrogen-bonded to Arg313 while His310 forms a hydrogen bond with Glu351 of helix α3 (Figure 3C). While two conformations of the sDMA binding pockect have been observed in the structure of SMN Tudor domain (32), it must be noted that the loop of P4_3_2_1_2 sDMA-BmPAPI is fashioned at the surface and interacting with crystal symmetry equivalents, raising the possibility that the observed conformational changes are influenced by crystal packing. Indeed, the loop in one of the two molecules in the asymmetric unit of the P4_3_2_1_2 *apo*-form is in the sDMA-bound conformation (the loop in the other molecule is partially disordered). Further studies, perhaps including structure determination of the full-length protein, are necessary to resolve whether sDMA binding influences the position of the loop or if the movement described here is merely a product of crystal packing.

### Comparison of BmPAPI and other Tudor domain-containing proteins

Tudor domains are conserved through many eukaryotic species. A search using BLAST shows that PAPI protein from *Drosophila melanogaster* (accession number NP_608657.1) shares 45% sequence identity with the extended Tudor domain of BmPAPI (33, 34). The top hits with known structure from human, mouse and *Drosophila melanogaster* are human Tdrd2 (PDB ID: 5J39, chain A), extended Tudor domain Td3 from mouse Tdrd1 (PDB ID: 4B9W, chain A), and eTud11 domain of *Drosophila melanogaster* Tudor protein (PDB ID: 3NTI), sharing a sequence identity of 41%, 31% and 34% with BmPAPI, respectively. Figure S1 shows the alignment of these four Tudor proteins by the COBALT multiple alignment tool (35). While the structure of *Drosophila* PAPI remains unknown, the structures of the other three Tudor domains, chain A of human Tdrd2, chain A of mouse Tdrd1 TD3 domain and chain A of *Drosophila* Tudor eTud11 domain, align well with the sDMA-bound BmPAPI structure, with RMSD values of 1.2, 1.9 and 2.2 Å for 153 residues aligned respectively, based on protein structure comparison service PDBeFold at European Bioinformatics Institute (Figure 4) (20, 22, 36–38).

**Figure 4.**
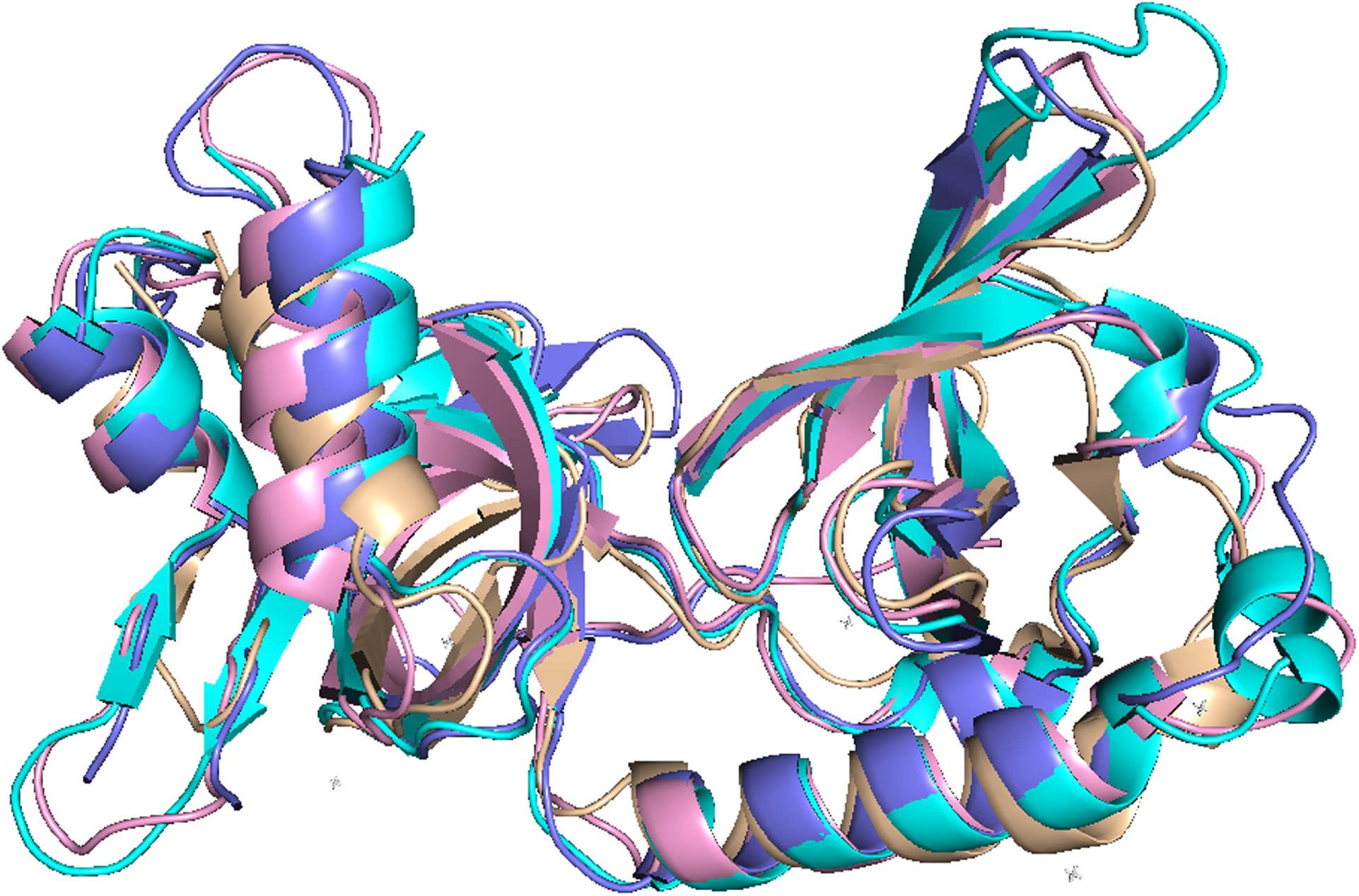
Superposition of the extended Tudor domain structures from BmPAPI homologs. Chain B of BmPAPI Tudor domain in the sDMA-bound state is shown in cyan; chain A of human Tdrd2 (PDB ID 5J39) is shown in lavender; chain A of Td3 domain from mouse Tdrd1 (PDB ID 4B9W) is shown in pink and eTud11 domain of *Drosophila melanogaster* Tudor protein (PDB ID 3NTI) is shown in wheat color.

### Aromatic cage of the core Tudor domain for sDMA binding

Similar to other PIWI-binding Tudor domains, four key residues of the BmPAPI core Tudor domain constitute the aromatic cage for sDMA binding: Phe308 in the loop region between β3 and β4, Tyr315 on β4, Tyr338 at the C-terminal end of β5, and Tyr341 on the loop linking β5 and β6 (Figure 5A). The four aromatic residues coordinate the guanidino group of sDMA in four orientations, with Tyr338 in the back, Tyr315 on the bottom, and Tyr341 and Phe308 sandwiching sDMA on each side. The plane of the guanidino group is approximately parallel to the aromatic rings of Phe308 and Tyr341. Figure 5B shows the negatively-charged surface of the aromatic cage area, which favors the binding of the positively-charged guanidino group of sDMA. All four residues form cation-π interactions with one of the two methylated amine groups, at distances of 3.7 Å, 4.1 Å, 5.3 Å and 3.7 Å between the nitrogen atom and the center of the benzene rings. The other methylated amine group is hydrogen-bonded to Asp343 on β6 to further enhance the binding interaction. A study on a homolog protein, *Drosophila* Tudor eTud11, identified a hydrogen bond between the amine group of sDMA and an asparagine residue in a similar position to BmPAPI Asp343 (22), and found that eliminating the hydrogen bond through mutagenesis caused a greater than 30-fold reduction of sDMA binding affinity compared to wild-type Tudor, indicating the critical role of this interaction in sDMA recognition. Although mutagenesis and affinity assays are yet to be conducted to confirm the role of BmPAPI Asp343 in sDMA binding, we expect it to be significant based on the high similarity between BmPAPI and *Drosophila* eTud11.

**Figure 5.**
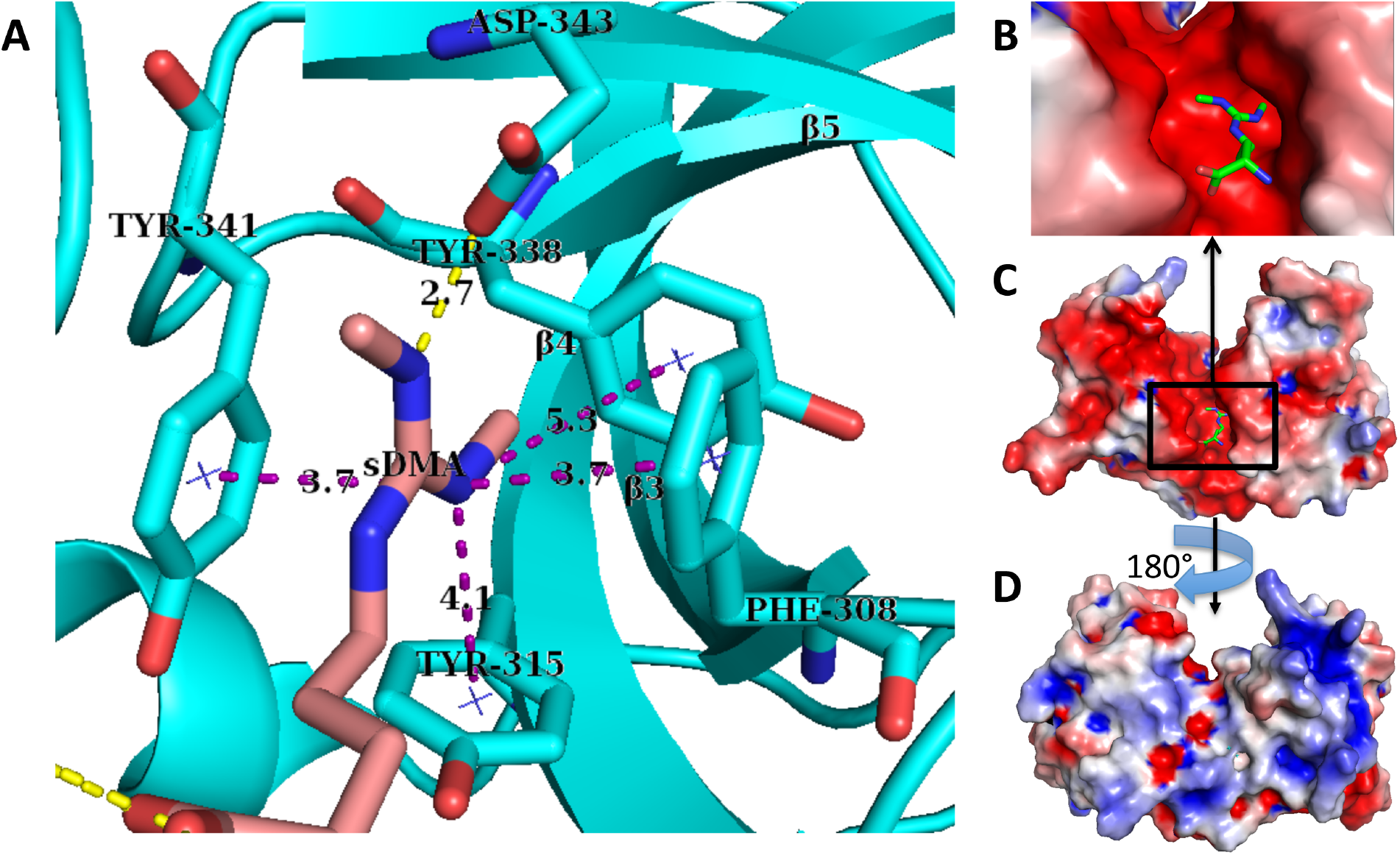
sDMA binding of BmPAPI. **(A)** A zoomed-in view of the aromatic cage for sDMA binding. The sDMA and interacting residues are shown in sticks with the sDMA colored pink. Blue and red indicate nitrogen and oxygen atoms respectively. Hydrogen bonds are shown as yellow dashed lines, while purple dashed lines show the distances between the nitrogen atom in the coordinated amine group and the centers of aromatic rings. **(B)** Surface electron potential of the aromatic cage, generated with PyMOL (The PyMOL Molecular Graphics System, Version 8, Schrödinger, LLC). sDMA is modeled in the bound position and shown in green sticks. The color of the surface reflects a range of −5.0 kT/e (red) to +5.0 kT/e (blue), colored by potential on the solvent-accessible surface. **(C, D)** Front and back view of the surface electron potential of BmPAPI extended Tudor domain. The color of the surface reflects a range of −3.0 kT/e (red) to +3.0 kT/e (blue), colored by potential on the solvent accessible surface.

### Surface potential of the extended Tudor domain of BmPAPI

At the C-terminus, BmPAPI contains two highly charged KH RNA-binding domains. To understand whether there is any compensatory charge distribution at the sDMA-binding domain, which might act to stabilize the protein, we examined the surface potential of the BmPAPI Tudor. The extended Tudor domain has opposing surface potentials on the sides proximal and distal to the sDMA binding site (Figure 5C). The surface of the side proximal to the sDMA binding site is mainly negatively charged (Figure 5D), whereas the majority of the surface on the distal side is positively charged, except for a few pockets. Considering that the N-terminus of BmPAPI contains a transmembrane domain preceding Tudor, we speculate that the surface distribution of Tudor might have a role in co-localization of BmPAPI and a PIWI protein on the outer surface of the mitochondria (28); the positively-charged distal side will favorably interact with anionic phospholipids of the membrane while the proximal side remains open for PIWI protein recruiting and docking. Validation for this role of the surface charge distribution is an ongoing investigation.

## SUMMARY

BmPAPI, a *Bombyx* homolog of the *Drosophila* PAPI protein, has been reported to play a crucial role in 3’-end formation of mature piRNAs. In this article we present the crystal structures of the extended Tudor domain of BmPAPI, both in its native state and in complex with sDMA. Since the crystals are in a different space group, the possibility of packing-induced conformational change could not be assessed. Nonetheless, a conformational change in the sDMA-binding site of a Tudor domain upon sDMA binding may be possible in the context of RNA binding to the two KH domains. In summary, the structural information herein reveals the molecular details of the interaction between the PAPI Tudor domain and sDMA, furthering understanding of the role of the PAPI-PIWI interaction in piRNA biogenesis pathways.

## AKNOWLEDGEMENT

This work was supported by the National Institutes of Health grant (GM106047, to YK/RM) and a Japan Society for the Promotion of Science Postdoctoral Fellowship for Research Abroad (to SH). We would like to thank technical support provided by Drs. Akashi Otaki and Shozo Honda in preparation of bacterial expression system.

