## Supplement Table S1 for "Structural studies of the Tudor domain from the *Bombyx* homolog of *Drosophila* PAPI: Implication to piRNA biogenesis"

### SUPPLEMENTARY MATERIAL

**Figure S1.** Sequence alignment of the extended Tudor domain from BmPAPI, Drosophila melanogaster PAPI (NP_608657.1), human Tdrd2 (5J39_A), mouse Tdrd1 (4B9W_A), and Tudor protein from Drosophila melanogaster (CAA44286.1).

```
BmPAPI     237   KYKRPETTGPSIEVYVSAVS-SPSRFWVQFVGPQV-AQLDDLVAHMTEYYSKKENREA[4]HVSVGQVVAAVFRHDGRWY   315
NP_608657  250   KLMASKGEGKPMEVYVSAVA-SPTKFWVQLIGPQS-KKLDSMVQEMTSYYSSAENRAK[4]APYVGQIVAAVFKFDEKWY   328
5J39_A     13    NLYFQ------GEVYVSASE-HPNHFWIQIVGSRS-LQLDKLVNEMTQHYENSVPEDL   TVHVGDIVAAPLPTNGSWY   81
4B9W_A     8     WTWVEFTVDETVDVVVCMMY-SPGEFYCHFLKDDAlEKLDDLNQSLADYCAQKPPNGF   KAEIGRPCCAFFSGDGNWY   83
CAA44286   405   ----SLTVGLTYDVVISYVEnGPYLFWVHLKSSDH--DLSTMMGQIERTKLKALAQAP   --ELGTACVARFSEDGHLY   473

BmPAPI     316   RARV[6]EFDSSQQVADVFYLDYGDSEYVATHELCELRADLLRLRFQAMECFLAGVRPAsgeEAVSPSGqtwDKWHPQAV   398
NP_608657  329   RAEI[6]QYNPKEQVIDLYFVDYGDSEYISPADICELRTDFLTLRFQAVECFLANVKSTiqtEPIT--------WPKSSI   403
5J39_A     82    RARV   LGTLENGNLDLYFVDFGDNGDCPLKDLRALRSDFLSLPFQAIECSLAR---------IAPSG---DQWEEEAL   146
4B9W_A     84    RALV   KEILPSGNVKVHFVDYGNVEEVTTDQLQAILPQFLLLPFQGMQCWLVD---------IQPPN---KHWTKEAT   148
CAA44286   474   RAMV   -CAVYAQRYRVVYVDYGNSELLSASDLFQIPPELLEIKPFAFRFALAGTKEI---EPIDDSMk--RIFKKSAI   544

BmPAPI     399   ERFEELTQVARWKALVSRTCTYK[2]ATAEGEKDKE---IPGIKLFDVTDEGELDVGAVLVAEGWAV----aGPAPSPRP   470
NP_608657  404   AKFEELTEVAHWRKLIARVVTYK[5]TTAVSAAAKEgtpLPGVELFDPADNSELNIADLMITQGFALPLddsYPVRSRSS   485
5J39_A     147   DEFDRLTHCADWKPLVAKISSYV   QTGIST-------WPKIYLYDTSNGKKLDIGLELVHKGYAIEL----PED----   208
4B9W_A     149   ARFQA---CVVGLKLQARVVEIT   ANGVG-----------VELTDLSTPYPKIISDVLIREQLVLRC----G------   201
CAA44286   545   YRNFELTVQAPESVGSMQTCHLN   QNGTNMLELLrqlKNSRQSYKKAEQLENDDAVEIRFIDSPSNF---YVQKVNI    618
```
